# *StrIPETrack*: a real-time, ROI-flexible tracking platform for high-throughput zebrafish behavior

**DOI:** 10.64898/2026.01.29.702662

**Authors:** Camden E. Cummings, Brandon L. Bastien, Jaqueline A. Martinez, Jinyue Luo, Summer B. Thyme

## Abstract

Quantitative phenotyping is essential to studies of animal behavior, enabling systematic analysis of variation arising from natural diversity or experimental manipulation. High-throughput behavioral assays that can simultaneously test multiple animals support sufficiently powered studies of behavioral variation, but accurate tracking of each animal is critical. Furthermore, behavioral tasks and experimental arenas span a wide range of complexity, from the reaction of a single larval zebrafish to an acoustic stimulus to associative conditioning in cue-rich environments. Here, we developed and validated *StrIPETrack* (Structural similarity-based Image Processing for Estimation and Tracking), a Python-based, modular animal tracking software designed for flexible region-of-interest (ROI) definitions and extensibility across assays. We show that *StrIPETrack* measures activity comparably to our previous LabVIEW-based zebrafish tracking software and detects similar behavioral differences between wild-type clutches. In addition, *StrIPETrack* accurately captures behavior in a complex arena: the Y-maze. Our approach for analyzing Y-maze navigation yields an expanded set of metrics beyond turn count and direction, revealing more subtle behavioral variation. Overall, this versatile software can be applied to monitor the activity of multiple animals in parallel in both simple high-throughput and more complex assays, and can be readily adapted to new paradigms.

**Summary:** Our open-source tracking software provides rich behavioral phenotyping of animals in many behavioral tasks. The flexible ROI design and live tracking makes the software adaptable to diverse paradigms.

## Introduction

Animal behavior arises from the coordination of multiple physiological processes, including nervous, endocrine, immune, muscular, and cardiac systems. Accordingly, variation in behavior can provide a sensitive functional readout of the effects of genetic and environmental perturbations on an animal. In model organisms, characterizing how genetic or neuronal perturbations alter behavior can also provide insights into the neurobiological basis of neurological and neuropsychiatric conditions. Therefore, reproducible and robust behavioral assays, coupled with sensitive quantitative analysis methods, are critical for both basic and translational animal research. High-throughput behavioral phenotyping in small model organisms enables the screening of hundreds of genetic mutations or thousands of small molecules for effects on behavior (MacRae and Peterson, 2015; Marcogliese et al., 2022; Yemini et al., 2013).

Zebrafish, particularly at the larval stage, have emerged as a powerful vertebrate model for high-throughput behavioral phenotyping due to their small size, rapid development, and relatively conserved brain architecture (Mueller and Wullimann, 2009). Genetic screens of more than one hundred mutants and pharmacological screens of over 10,000 compounds have been behaviorally characterized (Kokel et al., 2010; Thyme et al., 2019). Findings in zebrafish have also been translatable, as hits from zebrafish screens have entered clinical trials, such as for Dravet syndrome (Baraban et al., 2013; Patton et al., 2021). Previous scaling efforts have leveraged multi-well plates to measure sleep and circadian rhythms (Rihel et al., 2010), seizure-like behavior (Griffin et al., 2020), and responses to acoustic and visual stimuli (Randlett et al., 2019; Wolman et al., 2015).

Several software tools for real-time tracking of zebrafish exist. Bonsai (Lopes et al., 2015), an open-source visual programming language, and Zantiks, a proprietary tracking platform, both require adoption of software-specific programming environments. Our group’s previously published zebrafish tracking software (Joo et al., 2020) was implemented in LabVIEW, which requires a paid license. Open-source and Python-based alternatives exist (e.g., Stytra (Štih et al., 2019)), but in practice are often optimized for a particular arena geometry, animal number, hardware configuration, or stimulus delivery. A modular, open-source platform paired with an extensible behavioral apparatus (Joo et al., 2020) provides the greatest flexibility for innovative behavioral assay development.

In particular, scalable quantification of more complex behavioral tasks performed by juvenile zebrafish imposes software requirements that extend beyond those of commonly used larval assays. One such task is Y-maze navigation, which provides a measure of spatial working memory (Lalonde, 2002). One adaptation of this assay is the free movement pattern (FMP) Y-maze (Cleal et al., 2021a; Cleal et al., 2021b; Cleal et al., 2023). In this task, zebrafish freely explore an equilateral, three-armed Y-maze, and their pattern of turns is biased toward an alternation strategy (left-right-left-right/LRLR or right-left-right-left/RLRL) over other possible turn permutations. Pharmacological manipulation using drugs known to impair working memory disrupts performance of this assay in zebrafish (Cleal et al., 2021b). Notably, this bias is conserved across many vertebrate species, including humans (Cleal et al., 2021a).

To facilitate behavioral analysis across a wide array of zebrafish assays, we developed *StrIPETrack*, an open-source, Python-based tracking software. Our previously published zebrafish tracking system (Joo et al., 2020) was implemented in LabVIEW, which limits accessibility and is not easily extensible. *StrIPETrack* supports flexible ROI selection, enabling robust tracking of animals in simple and complex arenas. The positional information can be processed downstream or in real time, supporting quantification of multiple metrics, such as spatial preference within an arena and transitions between different zones. Here, we validate the tracking performance of *StrIPETrack* relative to our LabVIEW-based tracking in 96-well larval behavior assays and demonstrate its ability to accurately quantify turning behavior in the FMP Y-maze.

## Results

### Development and capabilities of *StrIPETrack*

Python-based *StrIPETrack* functions by having the user select regions of interest (ROIs) prior to the tracking process. Tracking is simplified by selecting the bounds of the areas where each fish will be. ROIs can be selected manually, through a graphical user interface (GUI) (**Fig. 1**). In *StrIPETrack*, each operation (i.e., ROI selection, tracking, y-maze analysis) is an entirely separate module, not requiring the use of others. This design decision means the software is both easy to add to and easy to incorporate into an existing pipeline. As previously (Joo et al., 2020), Python sends acoustic and visual stimuli commands to an Arduino and Teensy microcontroller.

**Fig. 1.**
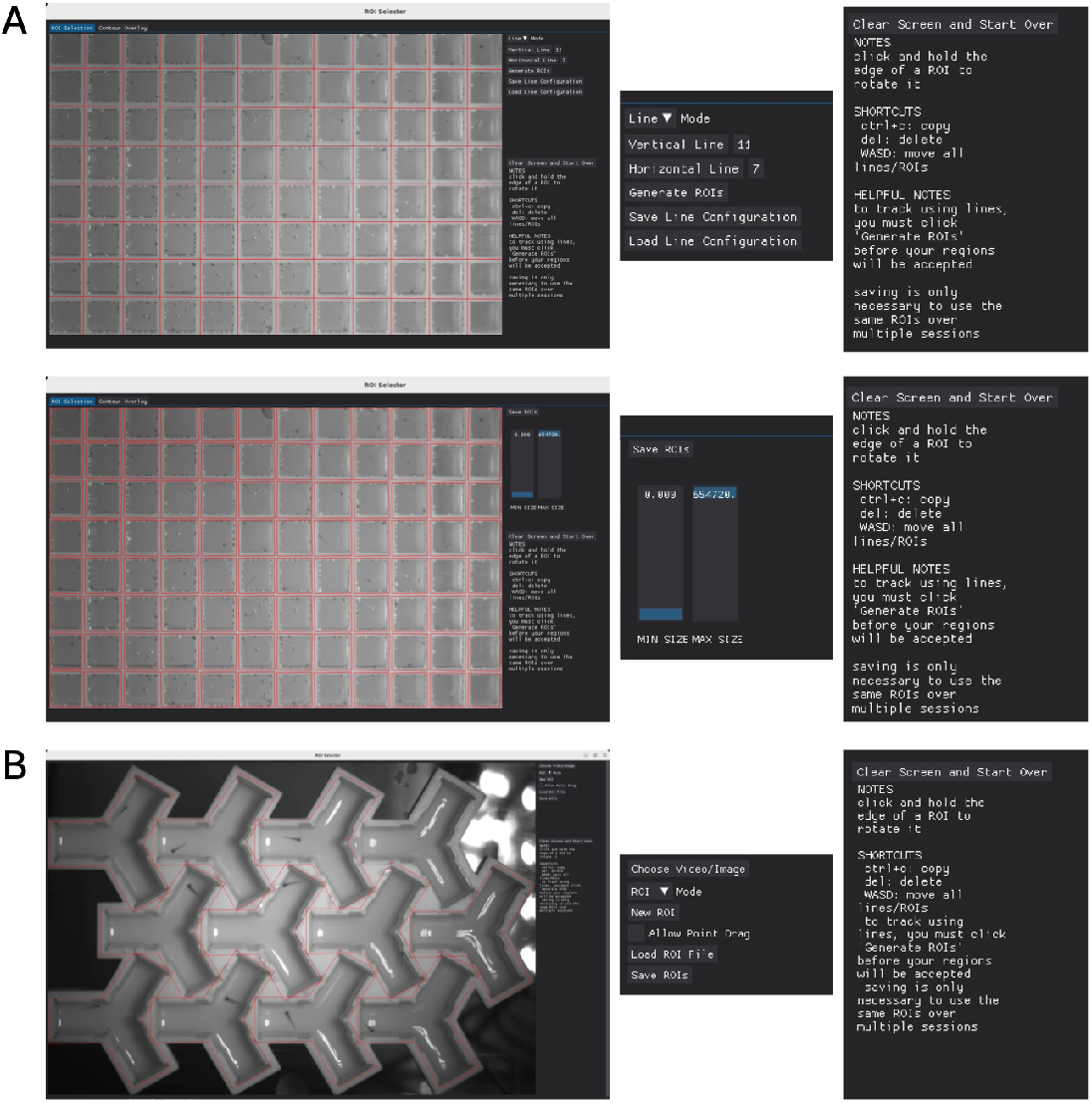
ROI preparation in *StrIPETrack*. Two ROI generation methods. The same set of functions are allowed in both. For example, keyboard shortcuts move lines/polygons, delete, and copy. Line or polygon files can be saved and re-opened and modified later. (A)The “Line” tool for convex ROIs. In “Line” mode, users can drag and drop lines or line edges. Then, to create ROIs bounded by each line, they can hit “Generate ROIs” – an easier option than manually creating each square using the “ROI” tool. (**B**) The “ROI” tool for concave ROIs.

For segmentation, we use structural similarity, an algorithm to assess the similarity of two images (Wang et al., 2004). Through this assessment, a difference image (i.e., a matrix of where difference is detected, **Fig. 2**) is created, which has been used repeatedly to identify animals for tracking (Chen et al., 2023; Loza et al., 2006). We use structural similarity to compare the current frame to a background frame, highlighting movement within each frame, as the change between a background frame and the current frame is the fish. To get an image containing only the background, we take a mode over the whole movie. If the fish is active, it will not be seen, and an image including only the background will be created. We use this image to segment the fish. Through optimization of the implementation provided by open-source library scikit-image (van der Walt et al., 2014), and altering of the algorithm for real-time tracking applications,significant speed improvements were achieved. Through this improvement, tracking at 30 fps is achieved on an Intel Core i7, 8GB RAM computer. Once the fish are segmented and a likely position is identified, we apply a Kalman Filter to smooth position data. Additionally, we discard tracks that do not move for many frames.

**Fig. 2.**
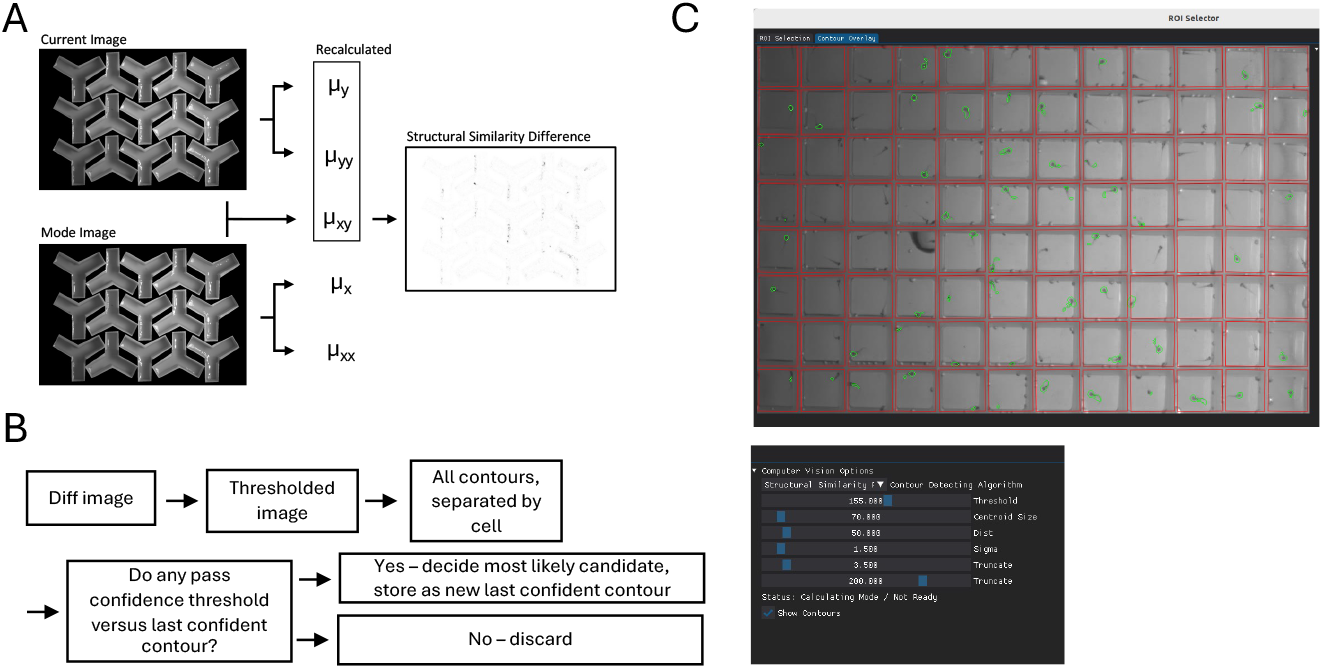
Structural similarity implementation. (**A**) In structural similarity, matrices µ _y_and µ_yy_ are calculated based on image 1, matrices µ_x_and µ_xx_are calculated based on image 2, and matrix µ_xy_ is calculated from both images. Because the sci-kit implementation of the algorithm does not anticipate a relationship between the two images, these values are recalculated each time. When the prior image is used, the current image’s µ_x_and µ_xx_ will become µ_y_and µ_yy_. When mode is image 1, µ_x_and µ_xx_ can be stored. (**B**) The difference image given by SSIM is thresholded, and contours for each cell are found. Each is checked for confidence, and if passes, is stored as new contour. (**C**) Detected zebrafish movements are highlighted in green, based on selection of the “Show Contours” checkbox in the GUI.

### Larval Behavioral Analysis Comparison to LabVIEW

First, we compared behavioral metrics generated from our existing LabVIEW-based tracking and our custom Python software. We previously demonstrated significant behavioral differences between larval clutches from different parents can be detected (Joo et al., 2020). To benchmark *StrIPETrack* against LabVIEW, we placed two different clutches of wild-type larval fish in alternating columns of a 96-well plate and tracked their movement with both software. Over a three-day time course with multiple stimuli (Capps et al., 2025; Joo et al., 2020), clear circadian-dependent activity differences were observed (**Fig. 3A, Fig. 3B**). Responses to dark flash were different between the two clutches. Specifically, pixel displacement (Δ pixels, dpix) in response to dark flashes was significantly higher in Clutch 1 than Clutch 2 in both methods (**Fig. 3C, Fig. 3D**). Thus, *StrIPETrack* captures similar behavioral variation comparable to LabVIEW, and is sufficiently sensitive to detect differences between groups of larval zebrafish.

**Fig. 3.**
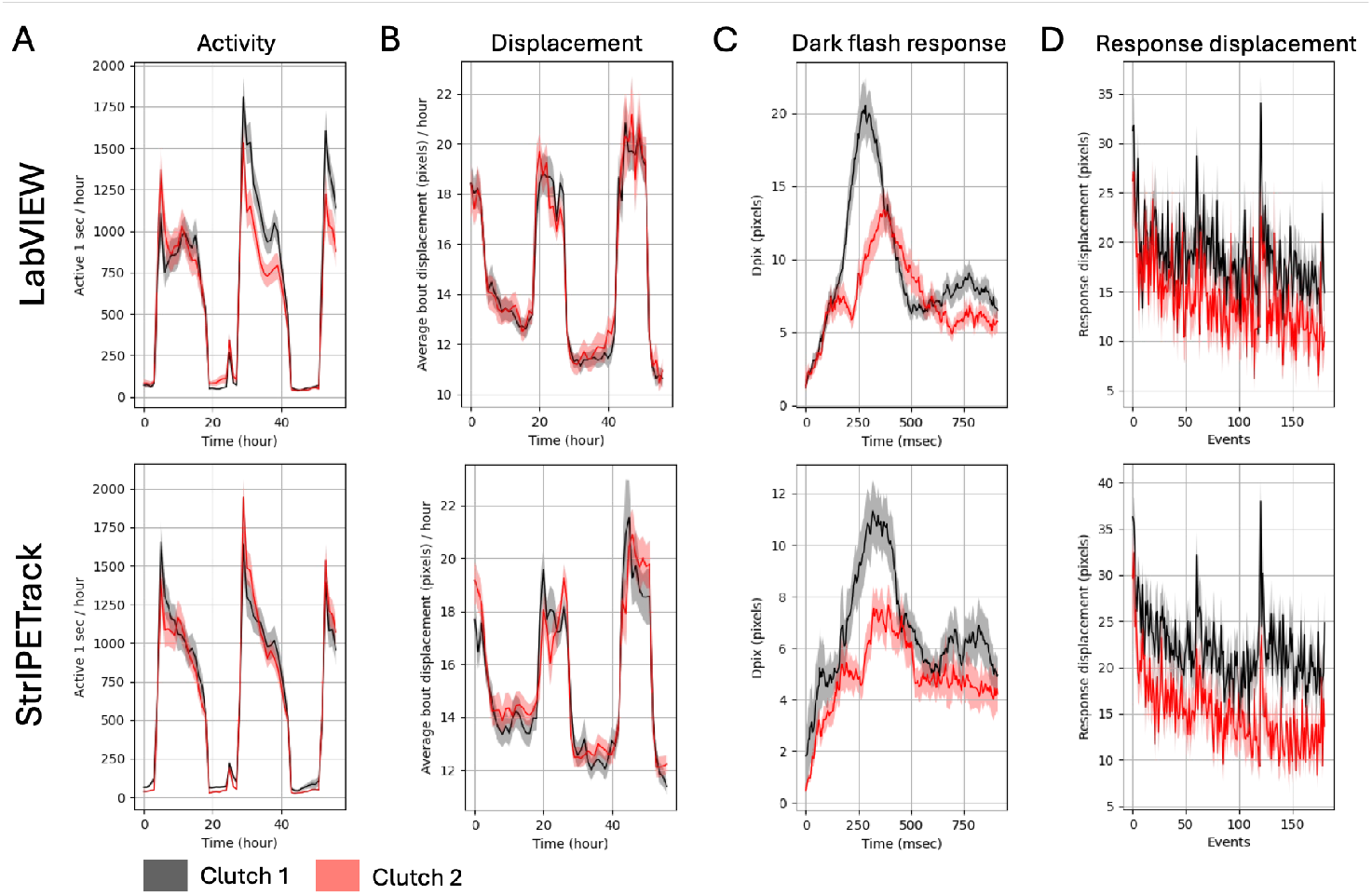
Comparison of 96-well plate larval behavior tracking between LabVIEW and *StrIPETrack*. (**A**) Active seconds per hours, a measure of overall activity. (**B**) Average bout displacement. This measure requires accurate tracking of the X-Y position. (**C**) Response trace for all dark flashes. Three blocks of 60 one-second-long dark flashes were delivered once per minute, with an hour break in between the blocks. Kruskal-Wallis Anova p-value LabVIEW = 0.0006, *StrIPETrack* = 0.001. (**D**) Response displacement for each dark flash. Kruskal-Wallis Anova p-value LabVIEW = 2.2e-5, *StrIPETrack* = 4.8e-8. LabVIEW clutch 1 N = 45, clutch 2 N = 43; *StrIPETrack* clutch 1 N = 41, clutch 2 N = 48.

### Free Movement Pattern (FMP) Y-maze Analysis Adaptability

To test the compatability of *StrIPETrack* with more complex ROIs, we used 3D-printed Y-mazes to adapt our behavioral apparatus to the previously published FMP Y-maze task. The common behavioral metric quantified in FMP Y-mazes is the turn direction into other arms (either left or right from the current arm) (Cleal et al., 2023). Sequences of four consecutive turns, called tetragrams, are calculated, and a preference for alternation is conserved across vertebrate species (Cleal et al., 2021a). To evaluate the accuracy of our code, we compared turns calculated by our software to turns measured by hand in a subset of fish at 10, 15, 21, and 27 days post-fertilization (dpf) using a Pearson correlation. There was a strong, significant positive correlation between the two methods (**Fig. 4A**). To assess if there was a systematic difference between our software and manual counting, we performed a paired t-test and found no difference between our software and manual (**Fig. 4B**). Thus, our code can accurately quantify tetragrams, highlighting its adaptability to complex ROIs.

**Fig. 4.**
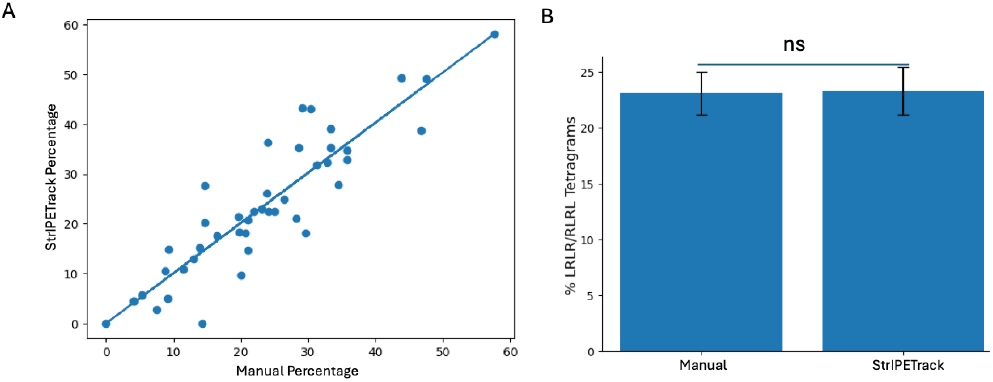
Validation of tetragram accuracy of Python tracking software. (**A**) Scatterplot plotting alternation tetragram (LRLR/RLRL) percent of individual fish measured by manual counting and our custom Python tracking software. Line represents the line of best fit (r=0.903, r^2^ = 0.82, p<0.0001, n=44). (**B**) Bar graph comparing the alternation tetragram percents for individual fish measured by manual counting and our custom Python tracking software. ns = not significant result of paired t-test: t=-0.22, p=0.82, n=44.

We next compared the performance of zebrafish in the FMP Y-maze across ages. When analyzing alternation tetragram percents, all ages preferenced alternation above random chance (**Fig. 5A**). To determine if performance in the task varied across time, we binned tetragrams performed into 10 minute bins. Zebrafish starting at 17 dpf performed alternating tetragrams in each bin above random chance. In contrast, the 10 dpf fish did trend toward alternations above random chance but were unable to reach False Discovery Rate-corrected in any of the ten-minute bins **(Fig. 5B, Fig. 5C)**. To assess for variability across wild-type fish, we compared alternation tetragram percentages across wild-type siblings from four different lines and found the LRLR/RLRL peak percentages ranged from as high as about 60% to as low as about 20% in 21 dpf fish. These percentages were always above random chance **(Fig. S2)**. We asked whether a simple biometric could predict performance in the task. We grouped the top 50% and lowest 50% of fish of 21 dpf fish based on body size and tested whether either group performed better or worse than the other. The top 50% of the fish performed the task better than the lowest 50% of fish, indicating that size could be a predictor of working memory performance performance (**Fig. S3**). We also stratified the 10 dpf fish into the upper and lower 50% of fish based on body size, and, like the 21 dpf fish, the bigger 10 dpf performed the task better than the smaller fish.

**Fig. 5.**
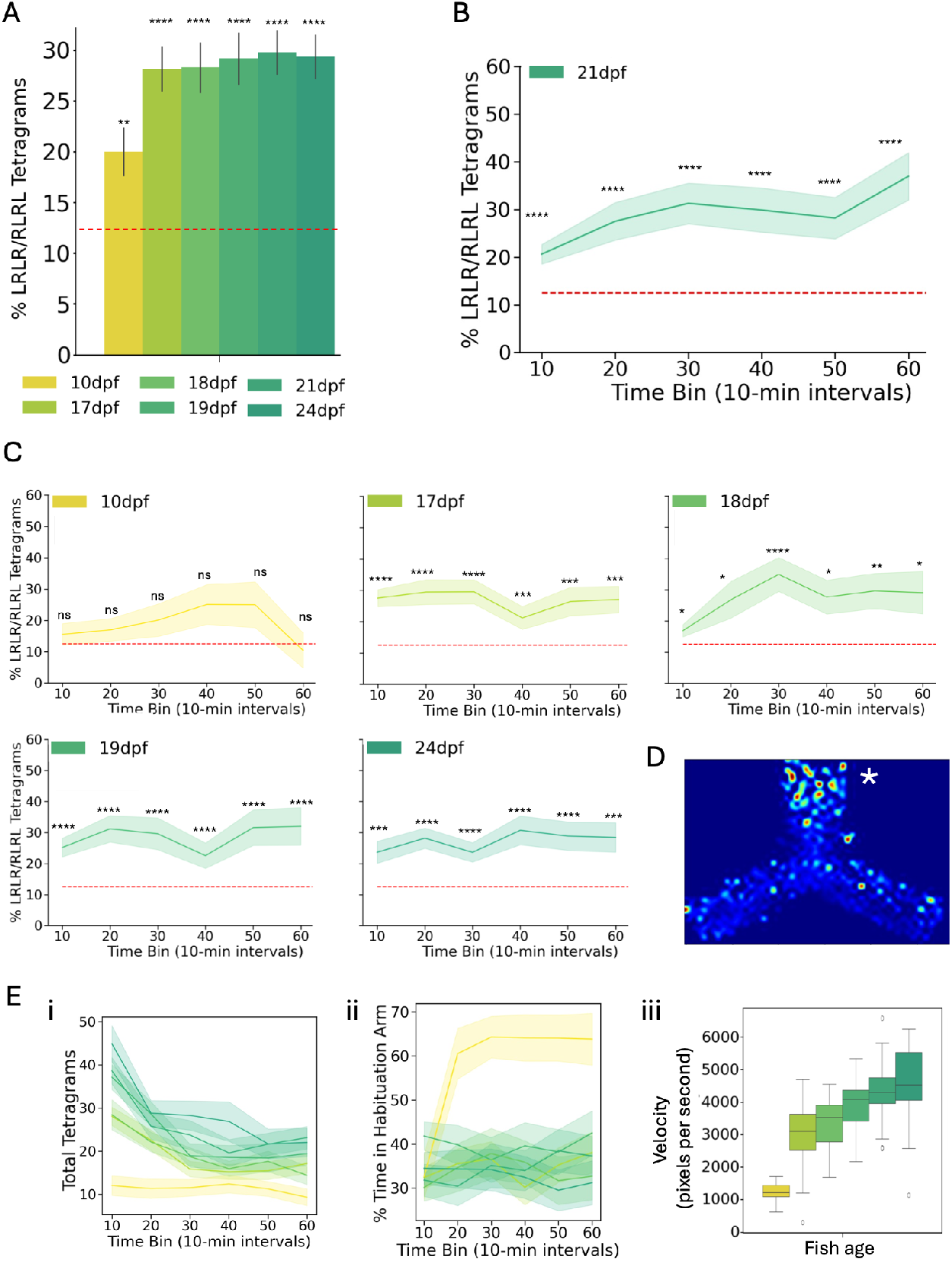
Behavioral metrics of wild-type fish of different ages. (**A**) Bar plot showing total alternation (LRLR/RLRL) tetragrams of wild-type zebrafish at different ages. (**B**) Examplar graph showing alternation tetragrams binned in 10-minute bins across the one hour recording. (**C**) Line graphs showing other ages alternation tetragrams in 10-minute bins. Statistical comparisons were made through one-sample t-tests against random chance for eight possible turn permutations (12.5%) ns=not significant, *<0.05, **<0.01, ***<0.001, ****<0.0001. (**D**) Heat map of localization in Y-maze. (**E**) Other behavioral metric, including i) total tegragrams, ii) time spent in habituation arm, iii) velocity. Error bars represent S.E.M..

Lastly, because our code was successfully able to quantify other behavioral metrics in larval behavior, we wanted to generate other behavioral metrics not previously quantified in Y-maze behavioral analysis pipelines. In addition to total tetragram numbers, which have been used in previous analyses, we quantified velocity, arm preference, and generated heatmaps representing position in the Y-maze (**Fig. 5D, Fig. 5E**). Total tetragrams and velocity provide readouts of overall activity, and the arm preference and heat maps provide readouts of arm bias, such as preference for the arm that the fish habituated in. These readouts supplement the tetragram percentage analysis by providing additional behavioral profiles and enhancing the interpretation of Y-maze behavior.

## Discussion

Here, we developed and validated an open-source, Python-based animal tracking software that is adaptable to simple and complex behavioral assays. Tracking larval zebrafish with *StrIPETrack* produced behavioral metrics comparable to those obtained with our established LabVIEW-based program (Joo et al., 2020), including the detection of significant differences between wild-type clutches (**Fig. 3D**). The magnitude of these differences reinforces the necessity of using sibling controls to achieve accurate results in zebrafish experiments. Leveraging the software’s ROI flexibility, we increased the capacity of the Y-maze assay to twelve arenas in a single plate that can be recorded simultaneously while being tracked independently. This scalability provides substantial potential for screen-based studies, such as investigating hundreds of mutant lines associated with neuropsychiatric disorders (Thyme et al., 2019) or screening compound libraries (MacRae and Peterson, 2015). Thus, this modular software supports standard larval behavior in 96-well plates and the generation of rich data sets for more complex paradigms.

Using our new analysis approach, we expanded the set of behavioral metrics extracted from Y-mazes, calculating velocity, preferences for zones, and activity (**Fig. 5D**). Previous analyses of the FMP Y-mazes mainly focused on turn-based metrics, including the number of turns and the percentage of tetragrams, reflecting the alternation strategy. While these analyses are critical, they may overlook more subtle behavioral features. The addition of the comb to habituate zebrafish is an effective strategy for synchronizing their navigation. With multiple custom-built behavioral arenas, more than one hundred animals can be assessed in a single day.

From validation of the Y-maze assay, we identified several variables that influenced performance. As previously reported (Cleal et al., 2023), animal age is a significant factor, with 10 dpf fish failing to perform tetragrams above random chance (**Fig. 5C**). We also noticed that body size affected performance at 21 dpf (**Fig. S3**), with larger fish performing better than smaller fish. Although causality is difficult to infer, improved working memory in larger fish could enhance hunting, allowing them to outcompete siblings. Arena size represents another variable that likely influences performance and should be scaled accordingly with age, particularly when testing fish older than 24 or younger than 10 dpf.

Beyond the assays tested here, *StrIPETrack* can facilitate other complex behavioral paradigms. For example, in social preference assays, the social stimulus fish is not always monitored (Capps et al., 2025; Dreosti et al., 2015; Geng et al., 2022), and our flexible ROI selection can easily track both fish. A particular strength of this tracking software is also the ability to track fish in real time. By tracking several fish in parallel in real-time, assays where fish automatically receive stimulation upon entering a region of an arena (e.g., a specific arm of a Y-maze) become possible using this software. This capability can allow for higher throughput testing of learning and memory-based tasks, such as aversive conditioning (Aoki et al., 2015).

Taken together, we have developed and validated *StrIPETrack*, a modular tracking software that provides expanded behavioral metrics in simple and complex tasks across a wide range of fish ages. Robust and reproducible behavior tracking is essential for identifying the underlying sources of variation in animal behavior. This software is highly adaptable to complex arenas and behavior analysis pipelines, representing a valuable resource for animal behavior studies.

## Materials and Methods

### Zebrafish Husbandry

Zebrafish experiments were approved by the UMass Chan Institutional Animal Care and Use Committee (IACUC protocol 202300000053). Animals were maintained on a 14 hr/10 hr light/dark cycle at 28°C. Larval fish were grown in 150 mm petri dishes until 5 dpf, or until used for larval behavior assays, at a density of less than 160 fish per dish. Debris was removed from the dish prior to 4 dpf. Only larvae that were healthy with developed swim bladders were used in this study. After 5 dpf, fish were maintained in 8700 ml tanks at a density lower than 50 fish/tank, unless otherwise noted.

### Larval Behavior Assays

Larval behavior assays were conducted as previously described (Capps et al., 2025; Joo et al., 2020). The custom-built behavior system consists of an infrared light source and camera overhead an acrylic platform. Larval zebrafish were placed in individual wells of a 96-well plate and sealed as previously described using a clear optical adhesive film (Thermo Fisher Scientific, 4311971). Plates were placed in the box in a 3D-printed fish plate holder. Our larval behavior pipeline includes dark flashes, light flashes, acoustic prepulse inhibition, and acoustic habituation. Acoustic and visual stimuli are controlled through a printed circuit board (PCB). Each experiment follows a file that dictates the times and durations of stimuli. We include a version of the PCB schematic from (Joo et al., 2020), which has been updated for Teensy 4.1, as the Teensy 3.6 used in the original paper is no longer available.

### Y-Maze Assays

To fabricate our Y-Mazes, we use the 3D-printed models shown in **Fig. 1B**, which are available on GitHub. We use the comb shown in **Fig. S1** to block the arms, so the fish habituate in one arm for 10 minutes prior to the beginning of the study. We print the Y-Mazes using white ABS, as it is both resilient and cost-effective. To avoid creating visual landmarks for the fish, the indentation for the comb is copied to each arm as well. We provide three versions of the comb: one for each arm. The indentation shown at the end of each Y-maze’s arm can be used to provide a stimulus for the fish, such as a coloured piece of plastic.

Y-maze assays were performed in the same behavior boxes as the larval behavior assays. A single zebrafish was placed in the individual arms of a Y-maze that was blocked off with a divider for ten minutes. Afterwards, the divider was lifted, and the fish was allowed to freely explore the Y-maze for one hour. Videos were recorded using the FLIR FlyCapture SDK software at 30 frames per second with H.264 compression using a streaming capture setting.

### Data Analysis

#### LabVIEW Analysis of Larval Behavior

We use the code provided in (Joo et al., 2020) as a comparison, as it demonstrates the expected behavior of *StrIPETrack*. It is a LabVIEW-based code for tracking, using the Visual Development Toolkit. It sends out stimuli at the appropriate times and tracks the delta pixels moved and the current position of each fish.

#### Manual Tracking of Y-maze tetragrams

To validate the accuracy of the software’s ability to detect entries into arms of the Y-mazes, we took a selection of random videos, spread across all ages of our data set, and manually tracked when fish entered and exited each arm. The data was analyzed to generate tetragrams, which were compared to the tetragrams generated by the tracking software.

## Acknowledgements

We thank the UMass Chan fish facility staff and the Research Computing team, as well as members of the Thyme lab and Phil Campbell for helpful feedback on the software.

## Funding

This research was funded by the following sources:

Simons Foundation SFARI Pilot Award (SBT)

National Institutes of Health [HD115159 to S.B.T., DP2NS132107 to S.B.T.]

## Data and resource availability

All data are available in the main text, the Supplementary Materials, or appropriate databases. Code is available from https://github.com/camden-cummings/zebrafish-tracker and https://github.com/camden-cummings/behavior-analysis-scripts. Files for generating 3D-printed arenas and updated PCB are available from https://github.com/camden-cummings/stripetrack_design_files.

## References

Aoki, R., Tsuboi, T. and Okamoto, H. (2015). Y-maze avoidance: an automated and rapid associative learning paradigm in zebrafish. Neurosci. Res. 91, 69–72.

Baraban, S. C., Dinday, M. T. and Hortopan, G. A. (2013). Drug screening in Scn1a zebrafish mutant identifies clemizole as a potential Dravet syndrome treatment. Nat. Commun. 4, 2410.

Capps, M. E. S., Moyer, A. J., Conklin, C. L., Martina, V., Torija-Olson, E. G., Klein, M. C., Gannaway, W. C., Calhoun, C. C. S., Vivian, M. D. and Thyme, S. B. (2025). Disrupted diencephalon development and neuropeptidergic pathways in zebrafish with autism-risk mutations. Proc Natl Acad Sci USA 122, e2402557122.

Chen, Z., Zhang, R., Fang, H.-S., Zhang, Y. E., Bal, A., Zhou, H., Rock, R. R., Padilla-Coreano, N., Keyes, L. R., Zhu, H., et al. (2023). AlphaTracker: a multi-animal tracking and behavioral analysis tool. Front. Behav. Neurosci. 17, 1111908.

Cleal, M., Fontana, B. D., Ranson, D. C., McBride, S. D., Swinny, J. D., Redhead, E. S. and Parker, M. O. (2021a). The Free-movement pattern Y-maze: A cross-species measure of working memory and executive function. Behav. Res. Methods 53, 536–557.

Cleal, M., Fontana, B. D., Double, M., Mezabrovschi, R., Parcell, L., Redhead, E. and Parker, M. O. (2021b). Dopaminergic modulation of working memory and cognitive flexibility in a zebrafish model of aging-related cognitive decline. Neurobiol. Aging 102, 1–16.

Cleal, M., Fontana, B. D., Hillman, C. and Parker, M. O. (2023). Ontogeny of working memory and behavioural flexibility in the free movement pattern (FMP) Y-maze in zebrafish. Behav. Processes 212, 104943.

Dreosti, E., Lopes, G., Kampff, A. R. and Wilson, S. W. (2015). Development of social behavior in young zebrafish. Front. Neural Circuits 9, 39.

Geng, Y., Zhang, T., Alonzo, I. G., Godar, S. C., Yates, C., Pluimer, B. R., Harrison, D. L., Nath, A. K., Yeh, J.-R. J., Drummond, I. A., et al. (2022). Top2a promotes the development of social behavior via PRC2 and H3K27me3. Sci. Adv. 8, eabm7069.

Griffin, A., Anvar, M., Hamling, K. and Baraban, S. C. (2020). Phenotype-Based Screening of Synthetic Cannabinoids in a Dravet Syndrome Zebrafish Model. Front. Pharmacol. 11, 464.

Joo, W., Vivian, M. D., Graham, B. J., Soucy, E. R. and Thyme, S. B. (2020). A Customizable Low-Cost System for Massively Parallel Zebrafish Behavioral Phenotyping. Front. Behav. Neurosci. 14, 606900.

Kokel, D., Bryan, J., Laggner, C., White, R., Cheung, C. Y. J., Mateus, R., Healey, D., Kim, S., Werdich, A. A., Haggarty, S. J., et al. (2010). Rapid behavior-based identification of neuroactive small molecules in the zebrafish. Nat. Chem. Biol. 6, 231–237.

Lalonde, R. (2002). The neurobiological basis of spontaneous alternation. Neurosci. Biobehav. Rev. 26, 91–104.

Lopes, G., Bonacchi, N., Frazão, J., Neto, J. P., Atallah, B. V., Soares, S., Moreira, L., Matias, S., Itskov, P. M., Correia, P. A., et al. (2015). Bonsai: an event-based framework for processing and controlling data streams. Front. Neuroinformatics 9, 7.

Loza, A., Mihaylova, L., Canagarajah, N. and Bull, D. (2006). Structural Similarity-Based Object Tracking in Video Sequences. In 2006 9th International Conference on Information Fusion, pp. 1–6. IEEE.

MacRae, C. A. and Peterson, R. T. (2015). Zebrafish as tools for drug discovery. Nat. Rev.Drug Discov. 14, 721–731.

Marcogliese, P. C., Deal, S. L., Andrews, J., Harnish, J. M., Bhavana, V. H., Graves, H. K., Jangam, S., Luo, X., Liu, N., Bei, D., et al. (2022). Drosophila functional screening of de novo variants in autism uncovers damaging variants and facilitates discovery of rare neurodevelopmental diseases. Cell Rep. 38, 110517.

Mueller, T. and Wullimann, M. F. (2009). An evolutionary interpretation of teleostean forebrain anatomy. Brain Behav. Evol. 74, 30–42.

Patton, E. E., Zon, L. I. and Langenau, D. M. (2021). Zebrafish disease models in drug discovery: from preclinical modelling to clinical trials. Nat. Rev. Drug Discov. 20, 611–628.

Randlett, O., Haesemeyer, M., Forkin, G., Shoenhard, H., Schier, A. F., Engert, F. and Granato, M. (2019). Distributed plasticity drives visual habituation learning in larval zebrafish. Curr. Biol. 29, 1337–1345.e4.

Rihel, J., Prober, D. A., Arvanites, A., Lam, K., Zimmerman, S., Jang, S., Haggarty, S. J., Kokel, D., Rubin, L. L., Peterson, R. T., et al. (2010). Zebrafish behavioral profiling links drugs to biological targets and rest/wake regulation. Science 327, 348–351.

Štih, V., Petrucco, L., Kist, A. M. and Portugues, R. (2019). Stytra: An open-source, integrated system for stimulation, tracking and closed-loop behavioral experiments. PLoS Comput. Biol. 15, e1006699.

Thyme, S. B., Pieper, L. M., Li, E. H., Pandey, S., Wang, Y., Morris, N. S., Sha, C., Choi, J. W., Herrera, K. J., Soucy, E. R., et al. (2019). Phenotypic Landscape of Schizophrenia-Associated Genes Defines Candidates and Their Shared Functions. Cell 177, 478–491.e20.

van der Walt, S., Schönberger, J. L., Nunez-Iglesias, J., Boulogne, F., Warner, J. D., Yager, N., Gouillart, E., Yu, T. and scikit-image contributors (2014). scikit-image: image processing in Python. PeerJ 2, e453.

Wang, Z., Bovik, A. C., Sheikh, H. R. and Simoncelli, E. P. (2004). Image quality assessment: from error visibility to structural similarity. IEEE Trans. Image Process. 13, 600–612.

Wolman, M. A., Jain, R. A., Marsden, K. C., Bell, H., Skinner, J., Hayer, K. E., Hogenesch, J. B. and Granato, M. (2015). A genome-wide screen identifies PAPP-AA-mediated IGFR signaling as a novel regulator of habituation learning. Neuron 85, 1200–1211.

Yemini, E., Jucikas, T., Grundy, L. J., Brown, A. E. X. and Schafer, W. R. (2013). A database of Caenorhabditis elegans behavioral phenotypes. Nat. Methods 10, 877–879.

