## Supplementary Information for "*StrIPETrack*: a real-time, ROI-flexible tracking platform for high-throughput zebrafish behavior"

### Contents


**1 Structure** **2**

**2 Real Time Tracking** **2**

**3 Tracking Videos** **2**

**4 ROI Selector DearPyGUI** **2**

**5 Y-Maze Analysis** **3**

**6 96-well plate Analysis** **3**

**7 Structural Similarity** **5**

**8 Supplementary Figures** **6**

### 1 Structure

To reduce complexity and increase modularity, the code has been divided into several Git submodules which can be used independently. Namely,

1. roi_selector_dearpygui - GUI for selecting desired ROIs
2. strsim_for_speed - structural similarity (computer vision method used)
3. y_maze_analysis - analysis of Y maze results & visualization

Our pipeline for real-time tracking is shown in the guitracker folder, and our pipeline for tracking videos is shown in the post file.

### 2 Real Time Tracking

For our tracking setup, we use a Grasshopper Gig-E camera. We access the camera using PySpin, a wrapper of Spinnaker. We provide both a GUI and a non-GUI interface for live-tracking, which are easily adapted to other cameras.

### 3 Tracking Videos

To track a folder of videos, simply open it using video-display-ex.py by inputting folder name into parameter '–folder'. This will open GUI, and you can save .cells files for each video in the folder. Next, run each video in the folder using str_sim_run.py. This will output a pre & post-processed csv containing position data [pre contains all candidates considered for fish, post is the final position], as well as csvs containing frame & detected arm, and frame & detected turn (L/R).

### 4 ROI Selector DearPyGUI

GENERAL FUNCTIONALITIES
**Delete:** To delete any object, simply press 'delete' while hovering over the object.
**Copy:** 'CTRL+C' copies the object that is being hovered over - they are not offset, so if you press 'CTRL+C' and nothing appears to happen, try to drag the object.
**Move All Objects:** Both ROIs and Lines can be shifted using the WASD keys, and every ROI/Line will be affected. This behavior is segregated s.t. if you are in ROI mode, it will only affect ROIs, and if you are in Line mode, it will only affect lines. W == UP, A == RIGHT, S == DOWN, D == LEFT
**Save:** Opens a directory tool, saves all current ROIs or lines to given filename.

LINE MODE
**Move Line:** To move any line, hover the mouse tool over the center point (or

either vertex), click, and drag.
**Generate ROIs:** Makes ROIs using lines as ROI bounds [see Fig.1]. After all ROIs are generated, all behaviors of ROIs still apply (i.e., you can still use WASD to move ROIs, 'del' to delete them, etc.)
**Clear All ROIs:** Deletes all current ROIs, returns to initial state.

ROI MODE
**Move ROI:** To move any ROI, simply hover the mouse tool over it, click, and drag - when you release the mouse tool, the ROI will be dropped in place.
**Rotate ROI:** To rotate any ROI, simply hover over any vertex of the ROI, click and drag.
**Drag ROI points:** To manipulate the size of the polygon itself (ex: if you made the ROI too small), click the " button. You are now in " mode, and can click and drag the vertices of any ROI to move it.

*What happens when I try to move an ROI out of bounds?*
The ROI will be placed back within bounds.

*Expanding or modifying:*
To change the appearance of the GUI itself, modify either video_gui [if you are attempting to change the video's visualization] or gui [to modify the GUI itself]. The relationship between the ROI Interface and Line interface, is mediated by state_manager.

#### 5 Y-Maze Analysis

To generate all papers listed in this paper, use paper_figures.ipynb or paper_figures.py.

*Arm Analysis*
Arm analysis takes in a csv with at minimum columns labeled 'pos_x', 'pos_y', 'frame', 'row' and 'col'. It outputs csvs with arm data, and tetragrams data.

#### 6 96-well plate Analysis

Parameters
-xhym - creates position based heatmaps
-e - same scheduled events used to schedule stimuli for run
-g - genotype ids, shown below
-s - sections file, shown below
-j - parameters for plotting
-r - cells file, generated by GUI
-m - path to experiment folder
-output_folder - output folder for data and graphs

-c - centroid file

-d - dpix file (LABVIEW only)
-t - timestamp file (LABVIEW only)

Python uses csv file containing positions, timestamps, and dpix, and inputs it here. LabVIEW splits those up into a postions file (.centroid), a dpix file (.motion) and a timestamp file (.timestamp).

| Parameter Name | Python | LabVIEW | Details |
|---|---|---|---|
| -xyhm | x | x | |
| -e | x | x | |
| -c | x | x | csv vs .centroid |
| -d | | x | .motion |
| -t | | x | .timestamp |
| -g | x | x | |
| -s | x | x | standard sections file in Git |
| -j | x | x | PlotParameters example in Git |
| -r | x | x | file ending in '.cells' vs a rois_string |
| -m | x | x | |

EXAMPLE PYTHON CALL:
python processmotiondata.py -python -xyhm -e /path/to/file/scheduled-events -c /path/to/file/pre-processed.csv -g /path/to/file/genotype_file -s /path/to/file/sectionsfile -j PlotParameters -r /path/to/file/zebrafish-tracker.cells -m /path/to/location/of/highspeedmovies/
EXAMPLE LABVIEW CALL:
python processmotiondata.py -xyhm -e /path/to/file/fulltestrun_final_01_27_2020 -c /path/to/file/'testlog.centroid1.Tue, Dec 23, 2025' -d /path/to/file/'testlog.motion1.Tue, Dec 23, 2025' -t /path/to/file/'testlog.timestamp1.Tue, Dec 23, 2025' -g /path/to/file/genotype_file -s /path/to/file/sectionsfile -j PlotParameters -r /path/to/file/rois_string -m /path/to/location/of/highspeedmovies/

*genotyping*

name_of_group:1,2,3,4 ...
other_name_of_group:10,11,12,13 ...

*sections file : pulls out specific times for analysis*

lightflash_day5dpflfday=1_9:10:00-1_9:26:00
ppi_day5dpfppi=1_9:37:00-1_15:00:00
habituation_day5dpfhab1pre=1_15:35:00-1_15:54:00
habituation_day5dpfhab1=1_16:03:00-1_16:05:00
...

#### 7 Structural Similarity

A simplified version of SSIM (assuming $\alpha = \beta = \gamma = 1$ and $C3 = \frac{C2}{2}$) can be implemented as such:

$\mu_x = mean(im_1)$
$\mu_{xx} = mean((im_1)^2)$
$\mu_{xy} = mean(im_1 im_2)$
$\mu_y = mean(im_2)$
$\mu_{yy} = mean((im_2)^2)$

$\sigma_x = (cov)(\mu_{xx} - \mu_x^2)$
$\sigma_y = (cov)(\mu_{yy} - \mu_y^2)$
$\sigma_{xy} = (cov)(\mu_{xy} - \mu_y \mu_x)$

$L =$ max data range (i.e. 255), $K1 = 0.01$, $K2 = 0.03$

$C1 = (K1 * L)^2$
$C2 = (K2 * L)^2$

$$diff = \frac{(2\mu_x\mu_y + C1)(2\sigma_{xy} + C2)}{(\mu_x^2 + \mu_y^2 + C1)(\sigma_x + \sigma_y + C2)}$$

When using SSIM for tracking on a video, you can improve speed by assuming one of two things - either you compare against a mode image or compare the previous image taken to the current image taken. Given this implementation, it's clear that $\mu_x$, $\mu_{xx}$, and $\sigma_x$ are all derived from im1 alone, meaning that if im1 is a mode image, they do not need to be recalculated. By the same trick - if you want to compare each previous frame to each current frame, taking im1 as the current image, $\mu_y -> \mu_x$, $\mu_{yy} -> \mu_{xx}$, and $\sigma_y -> \sigma_x$.

### 8 Supplementary Figures

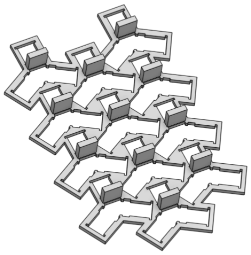

Fig. S1: Example y-maze comb for medium Y size. Fish are habituated in blocked arm, and after 10 minutes the comb is taken out and fish are released.

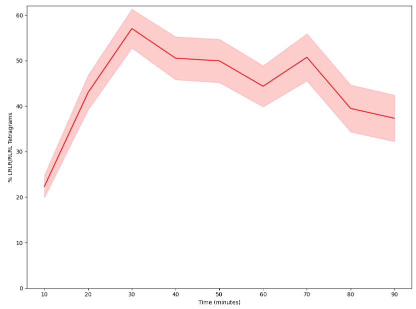


(a)

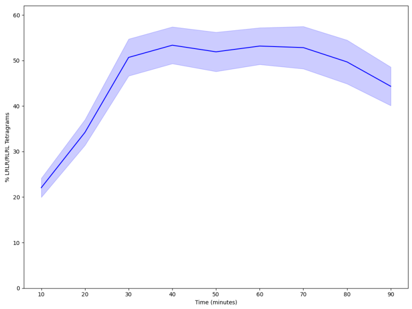


(b)

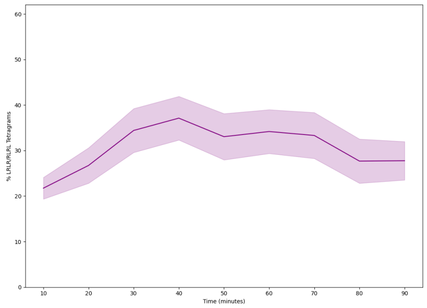


(c)

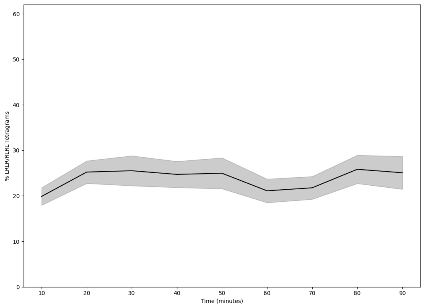


(d)

Fig. S2: Variation in wild type fish from different lines. Alternation tetragram percentage in 4 different lines of wild type fish over 90 minutes of tracking.

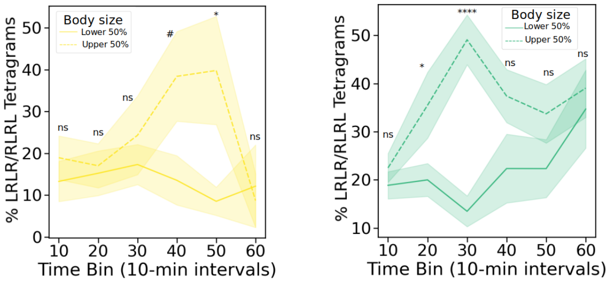


(a) (b)

Fig. S3: Body size influences performance in FMP Y-mazes. Comparison of alternation tetragrams percent of the largest 50% fish compared to the smallest 50% fish of 10 dpf fish (A) and 21 dpf fish (B). The results of two-way ANOVA are denoted above each time bin (ns = not significant, *<0.05, **<0.01. ***<0.001, ****<0.0001)
